# NCY-I β-lactamase activity correlates with antimicrobial susceptibility of a clinical strain of *Nocardia cyriacigeorgica*

**DOI:** 10.1101/2025.11.27.690932

**Authors:** Julie Couston, Athénaïs Le-Marchand, Elisabeth Hodille, Charlotte Genestet, Brune Joannard, Jérôme Feuillard, Yvonne Benito, Oana Dumitrescu, Mickaël Blaise

## Abstract

Nocardiosis is an infectious disease caused by several *Nocardia* species, among which *Nocardia cyriacigeorgica* is one of the most frequently isolated species in the clinic. Albeit most isolates of this species are susceptible to standard treatment combining trimethoprim and sulfamethoxazole, resistance has been reported, necessitating alternative or combination therapies. β-lactam antibiotics are of particular interest in this context. In this study, we aimed to address the β-lactam susceptibility profile of a clinical strain of *N. cyriacigeorgica* and assessed whether it correlated with the enzymatic activity of purified β-lactamase of the strain. We herein established that the strain is highly susceptible to imipenem and ceftriaxone, moderately sensitive to meropenem and resistant to amoxicillin. The resistance could be counteracted by β-lactamase inhibitors from two distinct chemical classes: vaborbactam, and avibactam while clavulanate was less potent. We demonstrated that the β-lactam susceptibility of the strain is in direct line with the enzymatic activity of purified NCY-I, a class A β-lactamase. NCY-I was indeed only active with amoxicillin but displayed poor activity towards other classes of β-lactams. The NCY-I activity could be inhibited *in vitro* by vaborbactam, clavulanate and avibactam. We consolidated these data by determining the high-resolution structure of NCY-I bound to avibactam. The structural analysis supported a conserved inhibitor binding site among other *Nocardia* class A β-lactamases strongly suggesting a broad inhibition spectrum of avibactam across *Nocardia* species.

## Introduction

*Nocardia* are environmental bacteria of the Corynebacterial/Mycobacterial order. To date, about 150 different and unique species of *Nocardia* have been reported (https://lpsn.dsmz.de/genus/nocardia). A third of these species are pathogenic for humans. Nocardia mediated infections are named nocardiosis. *Nocardia cyriacigeorgica* (*Ncyr*) is among the most frequently encountered *Nocardia* species in the clinic (Brown-Elliott et al., 2006; Conville et al., 2017). Recent genomic studies showed that *Ncyr* possesses the full arsenal of a true pathogen. The genome encodes virulence factors as well as elements essential for host cell invasion, intracellular survival and escape from cellular responses. Additionally, drug resistance genes have been reported (Yang et al., 2025).

Mild skin or lung nocardiosis is commonly treated with trimethoprim-sulfamethoxazole (TMP-SMX) monotherapy (Margalit et al., 2021; Yetmar et al., 2025a). In addition to first-line treatment and in cases of severe, disseminated infections and/or serious side effects to TMP-SMX, other classes of antibiotics can be given to treat nocardiosis. In this line, aminoglycosides can be considered, such as amikacin which is notably very potent against most nocardia species. Similarly, linezolid, a synthetic oxazolidinone, is very effective against all *Nocardia* species (Yetmar et al., 2025b).

Besides the standard treatment, penicillin derivatives are also of interest. However, β-lactam susceptibility may vary among *Nocardia* species (Hershko et al., 2023; Toyokawa et al., 2021). For instance, susceptibility to ceftriaxone, a third-generation cephalosporin, differs greatly, while *N. otitidiscaviarum* and *N. farcinica* are unsensitive, *N. brevicatena, N. paucivorans,* and the *N. abscessus* complex are highly susceptible (Hamdi et al., 2020). The presence of β-lactamases, known to hydrolyze β-lactams efficiently, should be considered in explaining such variations. So far, only class A β-lactamases have been reported in *Nocardia,* mainly using genomic or molecular biology methods.

Direct biochemical assays reporting β-lactamase activity were established for several reference strains of *N. farcinica*, *N. asteroides* and *N. brasiliensis* (Laurent et al., 1999; Lebeaux et al., 2020; Poirel et al., 2001; Songo et al., 2025). We recently reported the expression, and purification of a class A β-lactamase, NCY-I, from a *N. cyriacigeorgica* clinical strain as well as its crystal structure, representing the first one for this genus.

In this study, we report the β-lactam susceptibility profile of the above-mentioned *N*. *cyriacigeorgica* clinical strain as well as the potency of β-lactamase inhibitors in combination with β-lactams. We further characterized the enzymatic and inhibitory properties of the purified NCY-I. Finally, we report the crystal structure of NCY-I bound to avibactam at a very high resolution.

## Methods

### Determination of minimal inhibitory concentration

The MICs of four beta-lactams (amoxicillin, ceftriaxone, imipenem and meropenem (Sigma-Aldrich, Saint-Louis, MO, USA)), alone and in the presence of three beta-lactamase inhibitors (avibactam, clavulanate (Sigma-Aldrich, Saint-Louis, MO, USA), and vaborbactam (MedChemExpress, Vaulx-en-Velin, France) were determined for the clinical strain of *N. cyriacigeorgica* by 96-well broth microdilution as recommended (ISO, 2019). The 96-well plates were prepared with 50 µL of antibiotic solutions so that the final concentrations ranged from 0.12 μg.mL^-1^ to 128 μg.mL ^-1^ for beta-lactams with dilutions of 2 in 2, and fixed concentrations for inhibitors of 4 μg.mL^-1^ for avibactam, 2 μg.mL^-1^ for clavulanate and 8 μg.mL^-1^ for vaborbactam. For the inoculum preparation, *Nocardia* colonies cultured on Columbia agar with 5% sheep blood (COS, bioMérieux, Marcy l’Etoile, France) during 48 to 72h, in an aerobic atmosphere, were emulsified using a moistened swab in demineralized water, to obtain a bacterial suspension at 0.5 McFarland turbidity standard. Then 100 µL of the suspension was transferred into Muller-Hinton cation adjusted broth (Thermo Fischer Scientific, Waltham, MA, USA). The 96-well plate were inoculated with 50 µL of *N. cyriacigeorgica* inoculum. The plates were incubated at 35°C for 48h and the MICs were read visually using an inverted mirror. The determination of the MIC was carried out in three independent experiments. Quality controls using *Escherichia coli* ATCC 25922 and *Klebsiella pneumoniae* ATCC 700603 strains were performed to check the concentrations of the antibiotic solutions used.

### Expression and Purification of NCY-I

The cloning, expression and purification of NCY-I were essentially performed as described previously (Feuillard et al., 2024). Briefly, the protein lacking its secretion signal peptide and predicted unstructured part in the N-terminus, was expressed as a fusion with multi-histidine and SUMO tags in *E. coli* BL21. NCY-I was first purified on immobilized metal affinity chromatography (IMAC) (Ni-sepharose, Cytiva) after which both tags were cleaved by a His-tagged tobacco etch virus (TEV) protease. A second IMAC step was performed to get rid of the tags and the TEV protease, and the solution was further concentrated and loaded on a Superdex 75 Increase 10/300 GL size-exclusion chromatography (SEC) column (Cytiva). The last purification step on SEC was either performed using 20 mM Tris pH 8, 0.2 M NaCl, 0.5 mM β-mercaptoethanol and 5% glycerol or phosphate buffer saline (PBS) 1x and 10% glycerol as elution buffers, for structural studies or enzymatic assay purposes respectively.

### β-lactamase activity assays

The enzymatic assays (specific activity or kinetics) were made in PBS 1X buffer, and 1, 5 or 500 nM enzyme, depending on the substrate tested. The reaction was performed at 25℃ in a final volume of 100 μL. A calibration curve was established for each substrate to determine a correction factor and estimate product formation. For inhibition assays and IC_50_ determination NCY-I at a concentration of 8 nM, was incubated with the inhibitor at different concentrations for 15 min prior to mixing it in a 1:1 ratio with the solution containing nitrocefin at 400 μM. The reaction was then recorded for 30 min. Measurements were performed in 96-well plates (UV-star microplates, Greiner) and recorded on a Tecan Infinite 200 PRO M Plex plate reader. Kinetic constants were determined by plotting the data using the Michaelis-Menten curve function from GraphPad Prism v10. Catalytic efficiencies (*k*_cat_/*K*_M_) under substrate-limiting conditions were obtained from initial reaction rates and considering that *v≈ k*_cat_/*K*_M_*[E_t_], where *v* is the initial rate and [E_t_] represents the enzyme concentration.

### Crystallization, data collection and structure refinement

Crystals were obtained by mixing 1 μL of protein solution at a concentration of 14 mg.mL^-1^ with 1 μL of reservoir solution composed of 0.1 M citric acid pH 6.5, 6 % ethylene glycol and 10 % Polyethylene Glycol 6000. Ten days after setting up the crystallization experiment a 2 μl drop of solution, containing 0.1 M citric acid pH 6.5, 20 % ethylene glycol, 10 % Polyethylene Glycol 6000 and 10 mM avibactam, was added on top of the crystal drop and left overnight at 18°C. The crystals were then mounted in a litholoop and cryo-cooled in liquid nitrogen. Data collection was performed at the PXI-X06SA beamline at the Swiss Light Source, Paul Scherrer Institute. The structure was solved by molecular replacement using Phaser (McCoy, 2007) from the Phenix software suite (Liebschner et al., 2019) and the NCY-I structure (8qnn) used for phasing. The structure was manually adjusted with Coot (Casañal et al., 2020) and refined with the Phenix software suite. Structural data were deposited at the Protein Data Bank under the accession number: 9TEB.

## Results and discussion

We formerly reported the structural and preliminary biochemical characterization of NCY-I, a class A β-lactamase from a *N. cyriacigeorgica* clinical strain isolated from an elderly patient who was presenting a severe pulmonary infection (Feuillard et al., 2024). As this study was preliminary, we further wanted to investigate the β-lactam drug resistance profile of this strain and whether this correlated with the *in vitro* β-lactamase activity profile of the purified NCY-I. We first determined the drug susceptibility of the *N. cyriacigeorgica* clinical strain by testing three classes of β-lactams used at the clinic (**Table I**).

**Table I.** MIC (μg.mL^-1^) determination of β-lactams with or without β-lactamase inhibitors.

| β-lactams | No inhibitor | +Vaborbactam | +Clavulanate | +Avibactam |
| --- | --- | --- | --- | --- |
| **Amoxicillin** | 64-64-64 | 2-2-2 | 32-32-32 | 2-2-2 |
| **Ceftriaxone** | 4-2-2 | 2-2-2 | 2-2-2 | 2-2-2 |
| **Meropenem** | 8-8-8 | 4-4-4 | 4-4-4 | 4-4-4 |
| **Imipenem** | 1-1-0.5 | 0.5-0.5-0.5 | 0.5-0.5-0.5 | 0.5-0.5-0.5 |

The MIC values reported are from three independent experiments.

We determined minimum inhibitory concentrations (MIC) in liquid culture media for four β-lactams. The most active molecules were ceftriaxone, a 3^rd^ generation cephalosporin, and imipenem a carbapenem, with MIC of 4-2 and 0.5-1 μg.mL^-1^ respectively. Another carbapenem, namely meropenem, was less active with an MIC of 8 μg.mL^-1^. Amoxicillin of the aminopenicillin class was poorly active as its MIC was 64 μg.mL^-1^. These data and particularly the one for amoxicillin were suggestive of poor susceptibility and/or the existence of antimicrobial resistance. As class A β-lactamase activity was reported in *Nocardia* species (Laurent et al., 1999; Lebeaux et al., 2020; Poirel et al., 2001; Songo et al., 2025) and as we formerly demonstrated the existence of NCY-I in this strain, we tested three classes of β-lactamase inhibitors to address whether antimicrobial resistance could be circumvented. We combined these inhibitors with the formerly tested β-lactams to assess if we could circumvent the putative activity of the β-lactamase. To do so, vaborbactam, clavulanate and avibactam, from the boronic acid, β-lactam and diazabicyclooctane classes, respectively, were tested in combination with the four β-lactams. It is important to mention that the three inhibitors were tested at different but fixed concentrations and following the standard procedures (Eucast, 2025) which recommend the use of avibactam, clavulanate and vaborbactam in combination with β-lactams at 4, 2 and 8 μg.mL^-1^ respectively. Strikingly, a non-significant MIC shift (no reduction or only 1-fold MIC reduction) was observed for clavulanate in combination with the β-lactam antibiotics tested. However, a significant 32-fold (corresponding to 5 serial dilutions) MIC reduction was noticed for amoxicillin in combination with avibactam and vaborbactam, but no or only 1-fold MIC reduction (corresponding to a single serial dilution) shift was observed for the penems or ceftriaxone (**Table I**).

To strengthen these data, we characterized the kinetic properties of the recombinantly expressed and purified NCY-I for the four β-lactams tested. We first determined the specific activity of the enzyme in the presence of 250 μM of each substrate. We included as well nitrocefin, a chromogenic cephalosporin that is used as a standard substrate for β-lactamase assay (**Figure 1A**, **Table II**). Under these conditions NCY-I was the most active with nitrocefin and amoxicillin. NCY-I was however 11, 18 and 28 times less active with imipenem, ceftriaxone and meropenem respectively (**Figure 1A**, **Table II**). We next refined these first observations by determining the catalytic constants of NCY-I for these substrates when possible. We noticed that NCY-I was highly active with nitrocefin (*k*_cat_/*K*_M_= 1.65 × 10^6^ M^-1^.s^-1^) (**Figure 1B**, **Table II**). NCY-I was rather efficient at hydrolyzing amoxicillin (*k*_cat_/*K*_M_= 1.12 × 10^5^ M^-1^.s^-1^). NCY-I was nonetheless about 15 times less active with amoxicillin as compared with nitrocefin (**Figure 1C**, **Table II**), which is explained by a lower substrate affinity as the two enzymes have similar Vmax and *k*_cat_ values (**Table II**). Concerning the carbapenem class and as enzyme saturation was not reached, *k*_cat_ and *K*_M_ could not be directly determined. Therefore, catalytic efficiencies (*k*_cat_/*K*_M_) were inferred from the initial rate that was deduced from the linear part of the Michaelis-Menten curve (see Materials and Methods) (**Figure 1D, E and Table II**). The catalytic efficiency of NCY-I was slightly higher for meropenem than for imipenem but remained very low as compared to that for amoxicillin, which was approximately 500 times greater. Similarly, kinetic constants could not be determined for ceftriaxone, but NCY-I catalytic efficiency was estimated in the same way as for the two penems. NCY-I also exhibited very weak activity toward ceftriaxone, with a catalytic efficiency about 160 times lower than that of amoxicillin. (**Figure 1F and Table II**). These observations relate to formerly published data on another *Nocardia* BLA notably the recently described BRA-1 from the *N. brasiliensis* HUJEG-1 reference strain that was highly active with nitrocefin, active on amoxicillin and poorly active on penems (Songo et al., 2025). NCY-I is much more active than the second β-lactamase encoded by *N. brasiliensis* HUJEG-1 genome and named BRS-1 which had a very moderate activity with all the substates that were tested. However, NCY-I is much more active than the second β-lactamase encoded in the *N. brasiliensis* HUJEG-1 genome, BRS-1, which displayed a very moderate activity towards all the tested substates. Nonetheless, NCY-I is ten times less active than the *N. farcinica* β-lactamase FAR_IFM10152_ whose catalytic efficacy for nitrocefin was 1.5 × 10^7^ M^-1^s^-1^ and for amoxicillin, 2.2 × 10^6^ M^-1^s^-1^. FAR_IFM10152_ was also moderately active on imipenem and meropenem (Lebeaux et al., 2020) but slightly more than NCY-I.

**Figure 1.**
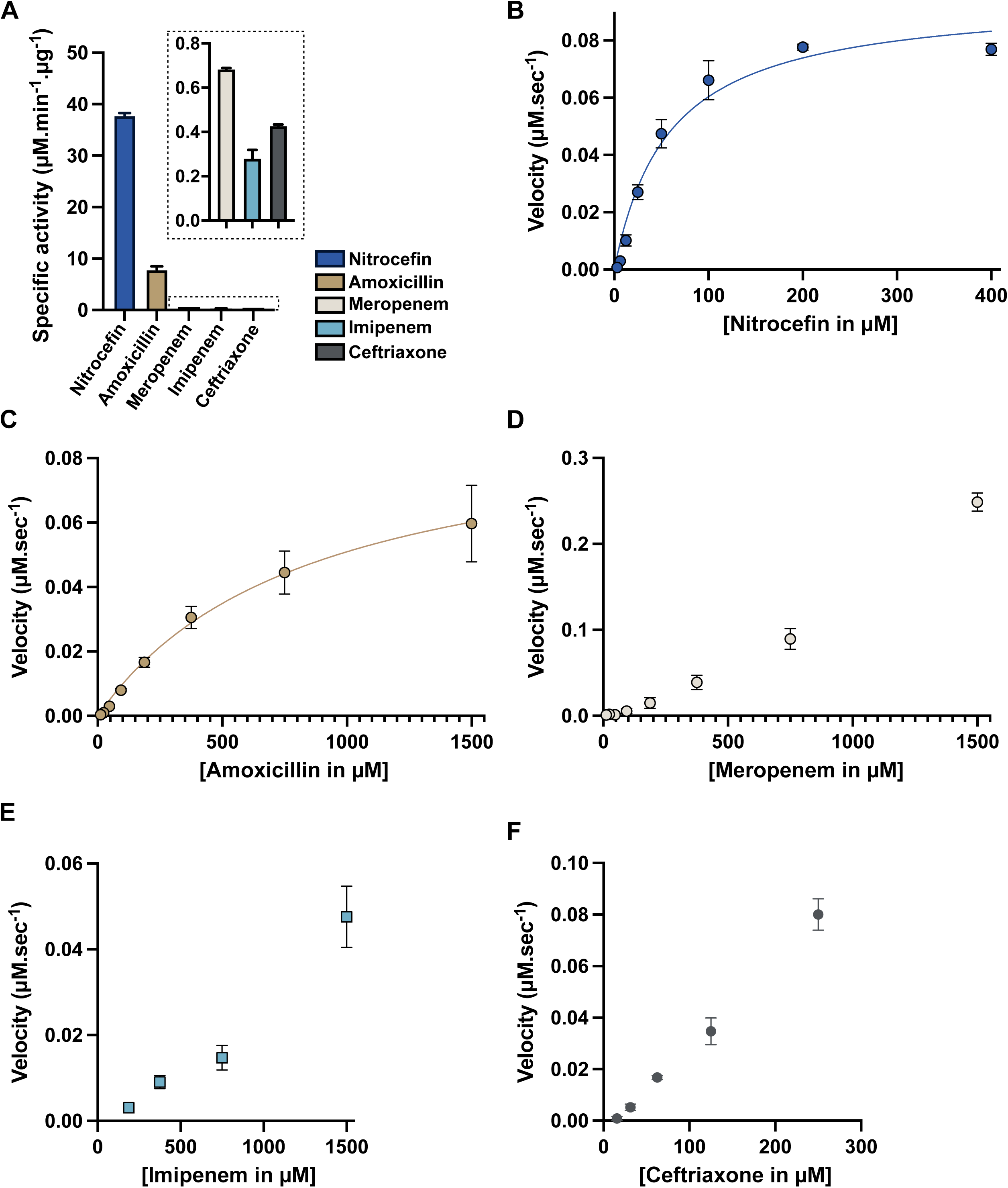
Enzymatic activity of NCY-I. A- Determination of the NCY-I specific activity. All substrates were tested at a concentration of 250 μM. Enzyme concentration was 1 nM for nitrocefin, 5 nM for amoxicillin and 500 nM for all the other substrates and the reaction recording was stopped at 30 min. B, C- Michaelis–Menten curves for reactions with nitrocefin and amoxicillin of NCY-I. D, E and F- Reaction rate for meropenem, imipenem and ceftriaxone. For all the graphs, data are the mean of three independent experiments and the error bars represent standard deviations.

**Table II.** Specific activity and kinetic constants of NCY-I for β-lactams.

| β-lactams | SA<br>($\mu\text{M}.\mu\text{g}^{-1}.\text{min}^{-1}$) | Vmax<br>($\mu\text{M}.\text{s}^{-1}$) | $K_M$<br>($\mu\text{M}$) | $k_{\text{cat}}$<br>($\text{s}^{-1}$) | $k_{\text{cat}}/K_M$<br>($\text{M}^{-1}.\text{s}^{-1}$) |
| --- | --- | --- | --- | --- | --- |
| **Nitrocefin** | $37.7 \pm 0.5$ | $0.095 \pm 0.003$ | $59.4 \pm 13.1$ | $95.7 \pm 3.8$ | $1.65 \pm 0.3 \times 10^6$ |
| **Amoxicillin** | $7.8 \pm 0.6$ | $0.097 \pm 0.03$ | $880.1 \pm 301$ | $97.2 \pm 29.5$ | $1.12 \pm 0.1 \times 10^5$ |
| **Ceftriaxone** | $0.42 \pm 0.007$ | nd | nd | nd | $6.83 \pm 0.6 \times 10^2$ |
| **Meropenem** | $0.68 \pm 0.008$ | nd | nd | nd | $2.34 \pm 0.3 \times 10^2$ |
| **Imipenem** | $0.28 \pm 0.04$ | nd | nd | nd | $0.66 \pm 0.1 \times 10^2$ |

We then investigated the difference of susceptibility of the *N. cyriacigeorgica* strain to β-lactamase inhibitors and assessed if this difference was also reflected on NCY-I *in vitro* activity. To perform the assay, we used nitrocefin as a substrate and titrated the three inhibitors. We next determined the IC_50_ values of three inhibitors, known to be suicide compounds inhibiting class A β-lactamase by covalent binding on the catalytic serine (Bush and Bradford, 2016). All three molecules inhibited 50% of NCY-I activity in the low μM range with IC_50_ for clavulanate, vaborbactam and avibactam of 7.2, 4.4 and 0.68 μM respectively (**Figure 2**). In agreement with our data from drug susceptibility testing, avibactam was the most potent in the enzyme inhibition assays.

**Figure 2.**
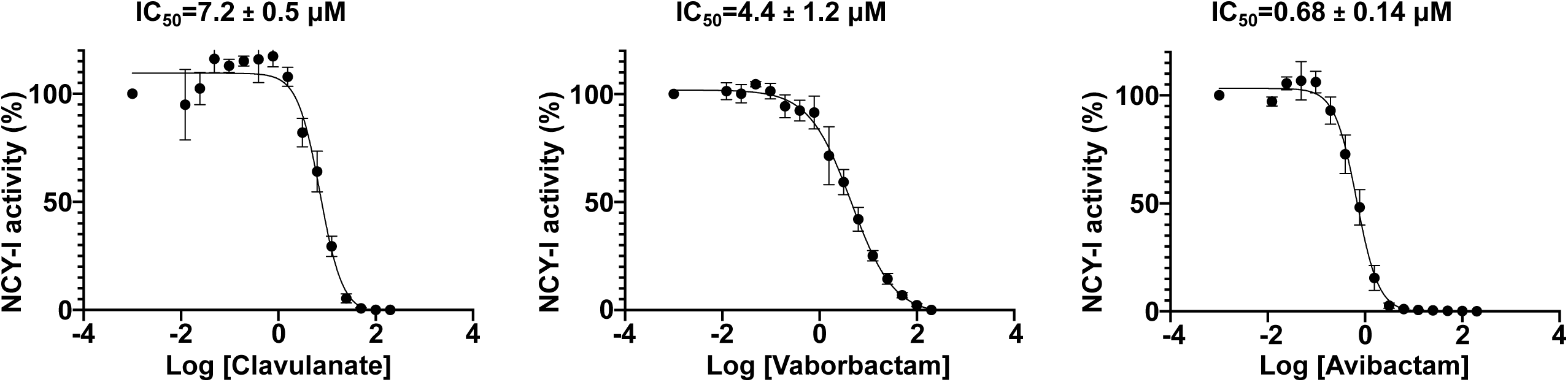
Inhibition of NCY-I. Dose-response curves of NCY-I inhibitors. Data are the mean of three independent experiments and the error bars indicate the standard deviation. The IC_50_ are fit of experimental data to the four-parameter sigmoidal dose–response equation of GraphPad Prism v10.

To complete our study, we attempted to solve the crystal structures of NCY-I bound to these three inhibitors but could only obtain the one with avibactam. The NCY-I avibactam complex structure was solved at very high resolution, 1.3 Å, thus enabling to model the structure with high accuracy (**Table III**). Unambiguously, the difference omit map demonstrated the presence of avibactam in the active site (**Figure 3A, B)**. As expected for this class of inhibitor, avibactam adopts an open structure and forms a covalent bond with the side chain of the catalytic Ser114, thus blocking the enzyme activity. Numerous residues are involved in the recognition of avibactam. Ser174, Asn176, Asn214, Thr260, Arg264, Thr279, Ser281 are involved in direct and water mediated H-bonds with the compound (**Figure 3C**). The tight binding is reinforced by a salt-bridge between the side chain of Lys278 and the sulfate group of avibactam. Interestingly, almost all these residues are very well conserved among the different β-lactamases retrieved from reference genomes, and particularly in the β-lactamases that were biochemically characterized and shown to be highly sensitive to avibactam (**Figure 4**). Indeed, sequence alignment of NCY-I, BRA-1, BRS-1 and FAR_IFM10152_ attests of a strong protein sequence conservation between the four enzymes as NCY-I displays 47% sequence identity with BRS-1, 64% with BRA-1 and 71% with FAR _IFM10152_.

**Figure 3.**
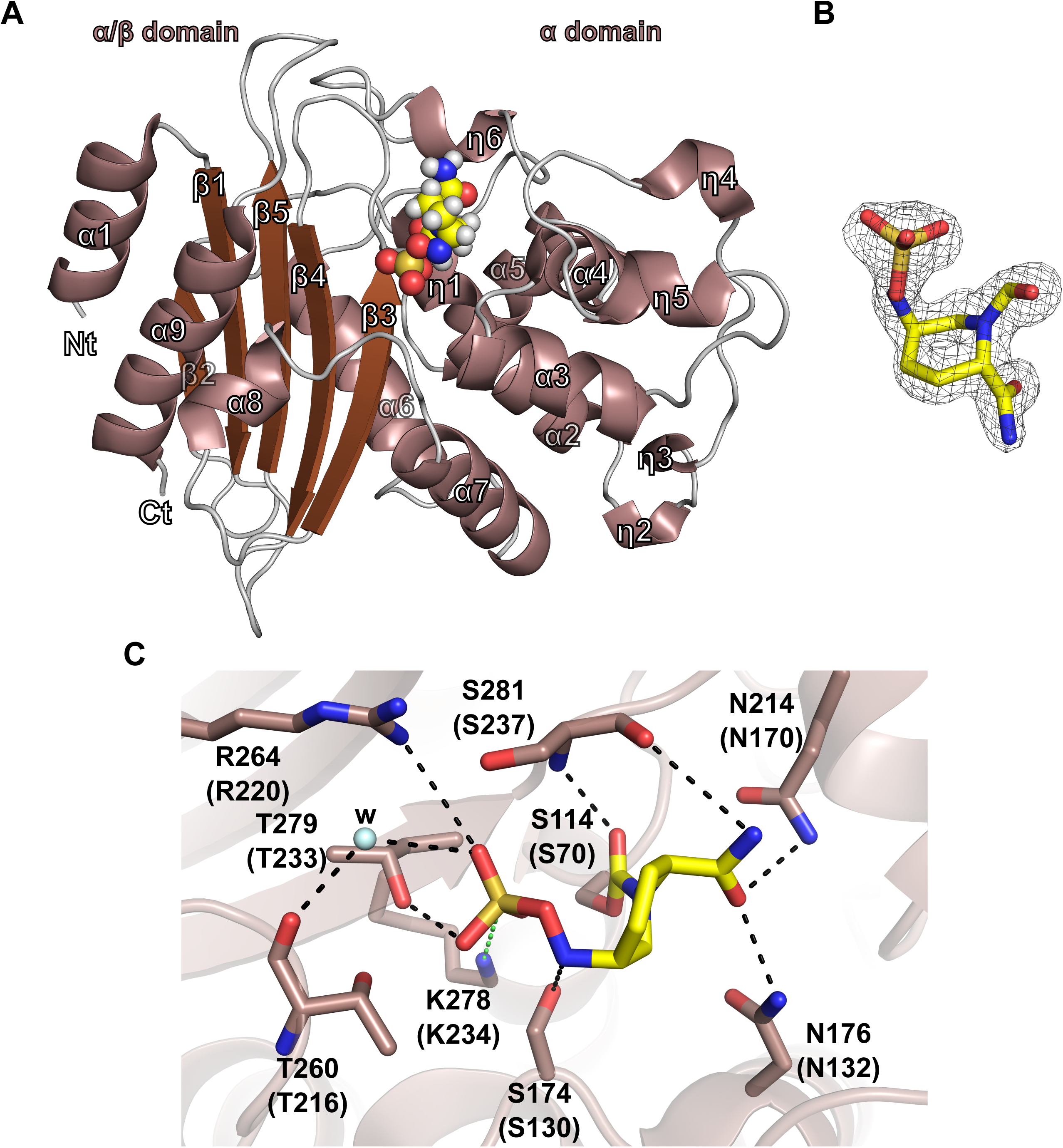
Structural analysis of NCY-1. A- Cartoon representation of the three-dimensional structure of NCY-1 bound to avibactam. α, β and η followed by a number indicate helices, strands and 3_10_ helices. Nt and Ct stand for N-terminus and C-terminus, respectively. B- Simulated annealing omit map supporting the presence of avibactam, shown as a yellow stick. The map is contoured at a level of 3.5 and represented as a black mesh. C- Depiction of the avibactam binding site. Residues interacting with avibactam are shown as sticks. W indicates a water molecule which is displayed as a light blue sphere. Black-dashed lines indicate hydrogen bonds and the green one represents a salt-bridge. Residues numbering corresponds to the protein sequence of NCY-I, while those in parentheses indicate Ambler numbering.

**Figure 4.**
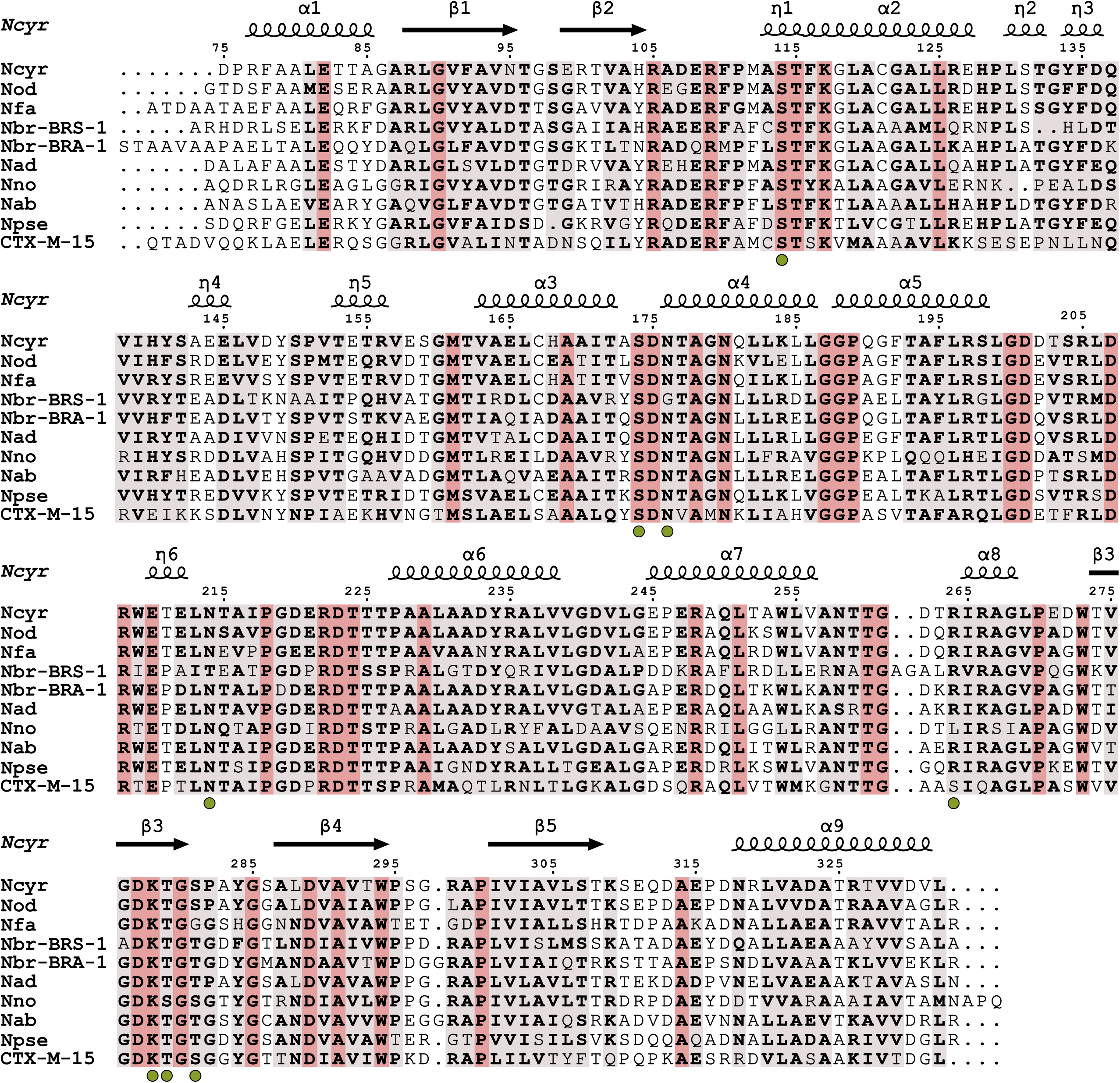
Multiple sequence alignment of class A β-lactamases from human pathogenic *Nocardia*. The sequence of NCY-I was aligned with the sequences of class A β-lactamases from *Nocardia otitidiscaviarum* NEB252, *Nocardia farcinica* IFM 10152, *Nocardia brasiliensis* HUJEG-1 (BRS-1 and BRA-1), *Nocardia asteroides* NCTC11293, *Nocardia nova* SH22a, *Nocardia pseudobrasiliensis* ATCC 51512, *Nocardia abscessus* NBRC 100374. CTX-M-15 was added as it is a well-studied and a model class A β-lactamase but not belonging to the *Nocardia* genus. Strictly-conserved residues are in red while semi-conserved ones are in light red. The green spheres below the sequences indicate the residues involved in avibactam recognition. The alignment displays only the sequence matching the NCY-I residues that were modeled in the structure. The secondary structure elements of the NCY-I structure are placed above the sequence alignment where α, β, η indicates α-helix, β-sheet and 3_10_ helix respectively. The figure was generated with the ENDscript server (Robert and Gouet, 2014) and manually adjusted in Inkscape.

**Table III.** Data collection and Structure refinement.

|  | NCY-I bound to Avibactam |
| --- | --- |
| **PDB accession number** | 9TEB |
| **Beamline** | PXI-X06SA, SLS |
| **Wavelength** | 0.999 |
| **Resolution range** | 44.55 - 1.3 (1.35 - 1.3) |
| **Space group** | *P* 3<sub>2</sub> 2 1 |
| **Unit cell (Å, °)** | 54.59 54.59 133.21<br>90 90 120 |
| **Total reflections** | 612900 (59564) |
| **Unique reflections** | 57487(6039) |
| **Multiplicity** | 10.6 (9.8) |
| **Completeness (%)** | 100 (100) |
| **Mean I/sigma(I)** | 16.7 (1.37) |
| **Wilson B-factor (Å<sup>2</sup>)** | 18.29 |
| **R-meas** | 0.064 (2.09) |
| **Reflections used in refinement** | 57484 (2678) |
| **Reflections used for R-free** | 2874 (134) |
| **R-work** | 0.156 (0.372) |
| **R-free** | 0.175 (0.352) |
| **Number of non-hydrogen atoms**<br>**macromolecules, ligands, solvent** | 2247<br>1996, 30, 221 |
| **Protein residues** | 259 |
| **RMS bonds (Å)** | 0.005 |
| **RMS angles (°)** | 0.74 |
| **Ramachandran (%)**<br>**favored, allowed, outliers** | 97.28, 2.72, 0 |
| **Clashscore** | 1.76 |
| **Average B-factor (Å<sup>2</sup>)**<br>**macromolecules, ligands, solvent** | 24.15<br>23.08, 23.37, 33.89 |

## Conclusion

To summarize, we investigated in this study the drug-susceptibility profile to β-lactams of a *N. cyriacigeorgica* clinical strain. We showed that this susceptibility pattern is in direct correlation with the *in vitro* activity of the purified NCY-I β-lactamase encoded by this strain. These observations suggest that no other β-lactamase activity is present in the strain and or that if any other β-lactamase is present it/they display(s) a similar activity to the NCY-I one. These results may suggest as well that there is no other antimicrobial resistance mechanism against these two β-lactams (amoxicillin and to a less extent meropenem) than the one mediated by β-lactamases. Interestingly, our structural study indicates that avibactam is tightly bound by residues that are highly conserved among β-lactamase from other *Nocardia* species thus strongly suggesting a broad spectrum of the inhibitor and as demonstrated by recent studies (Songo et al., 2025). Finally, the differences of the β-lactamase inhibitors efficacy against the strain can be partly explained by the difference in inhibition seen in the enzymatic assay and may reflect some mechanisms of resistance and/or poor permeability to clavulanate that awaits further investigation.

## Data availability

The data that support the findings of this study are available from the corresponding authors upon reasonable request.

## CRediT

**Julie Couston:** Conceptualization, Methodology, Formal analysis, Investigation, Writing – review and editing. **Athenaïs Le-Marchand:** Methodology, Formal analysis, Writing – review and editing. **Elisabeth Hodille:** Conceptualization, Methodology, Formal analysis, Supervision, Writing – review and editing. **Charlotte Genestet:** Methodology, Writing – review and editing. **Brune Joannard:** Supervision, Writing – review and editing. **Jérôme Feuillard:** Formal analysis, Investigation, Writing – review and editing. **Oana Dumitrescu:** Conceptualization, Methodology, Formal analysis, Resources, Project administration, Supervision, Writing – review and editing, Funding acquisition. **Mickaël Blaise:** Conceptualization, Methodology, Formal analysis, Investigation, Resources, Project administration, Supervision, Writing – original draft, Writing – review and editing, Funding acquisition.

## Acknowledgments

The authors acknowledge the ANR (French National Research Agency) for its financial support of the NORESIST project ANR-24-CE44-5883 attributed to MB and OD. We thank the Paul Scherrer Institute, Villigen, Switzerland for the provision of synchrotron radiation beamtime at beamline PXI-X06SA of the SLS. We are grateful to the CNRS, the Montpellier University and the IRIM directory for support.

## Ethics statement

None required

